# Fmo5 plays a sex-specific role in goblet cell maturation and mucus barrier formation

**DOI:** 10.1101/2024.04.05.588360

**Authors:** Megan L. Schaller, Madeline M. Sykes, Joy Mecano, Sumeet Solanki, Wesley Huang, Ryan J. Rebernick, Safa Beydoun, Emily Wang, Amara Bugarin-Lapuz, Yatrik M. Shah, Scott F. Leiser

**Author notes:** denotes co-corresponding authors.

## Abstract

**Background and Aims:** The intestine plays a key role in metabolism, nutrient and water absorption, and provides both physical and immunological defense against dietary and luminal antigens. The protective mucosal lining in the intestine is a critical component of intestinal barrier that when compromised, can lead to increased permeability, a defining characteristic of inflammatory bowel disease (IBD), among other intestinal diseases. Here, we define a new role for the flavin-containing monooxygenase (FMO) family of enzymes in maintaining a healthy intestinal epithelium.

**Methods:** Using *Caenorhabditis elegans* we measure intestinal barrier function, actin expression, and intestinal damage response. In mice, we utilize an intestine-specific, tamoxifen-inducible knockout model of the mammalian homolog of *Cefmo-2*, Fmo5, and assess histology, mucus barrier thickness, and goblet cell physiology. We also treat mice with the ER chaperone Tauroursodeoxycholic acid (TUDCA).

**Results:** In nematodes, we find *Cefmo-2* is necessary and sufficient for intestinal barrier function, intestinal actin expression, and is induced by intestinal damage. In mice, we find striking changes to the intestine within two weeks following Fmo5 disruption. Alterations include sex-dependent changes in colon epithelial histology, goblet cell localization, and mucus barrier formation. These changes are significantly more severe in female mice, mirroring differences observed in IBD patients. Furthermore, we find increased protein folding stress in Fmo5 knockout animals and successfully rescue the severe female phenotype with addition of a chemical ER chaperone.

**Conclusions:** Together, our results identify a highly conserved and novel role for Fmo5 in the mammalian intestine and support a key role for Fmo5 in maintenance of ER/protein homeostasis and proper mucus barrier formation.

## INTRODUCTION

The intestinal lining forms a tight barrier to dietary antigens and luminal microbes while also taking up nutrients, electrolytes, and water. The colonic epithelium is protected by a bi-layer of mucus directly adjacent to epithelial cells. The mucus barrier is composed of a dense inner layer with antimicrobial properties and a thick outer layer that is exposed to luminal contents^1^. Mucus maintains the intestinal barrier and is made solely by cup-shaped goblet cells (GCs) abundant in the colonic epithelium. Goblet cells constitutively synthesize mucins that, when secreted from storage vesicles, undergo biochemically-induced structural changes to expand, hydrate, and form an interconnected structure that makes up the inner layer of the colonic mucosal lining^2–4^. The absence of Muc2, the primary mucin in colonic goblet cells, leads to altered crypt morphology and impairments in goblet cell maturation and migration up the crypt^5^. These changes are detrimental to intestinal homeostasis.

The continuous requirement for colonic mucus production to maintain the barrier underscores the importance of the endoplasmic reticulum (ER) and unfolded protein response (UPR) in GCs. The ER is required to efficiently fold the large MUC2 protein but also with promptly detecting and managing misfolded proteins. Impairments in goblet cell ER function compromises the mucus barrier. Dysfunction in mucin processing and exocytosis from GCs is observed in patients with IBD (ulcerative colitis (UC))^6^, negatively impacting mucus barrier function. Additionally, in mice with colitis and humans with UC, the inner mucus layer is infiltrated with microbes, suggesting a failure in the antimicrobial function of the mucus barrier^7^. This increased permeability can result in inflammation, alterations in intestinal absorption, and damage to the intestinal epithelium^7,8^.

Flavin-containing monooxygenases (FMOs) are ER-resident enzymes that play an important role in xenobiotic metabolism^9,10^. In previous studies, we discovered that *fmo-2* in *C. elegans* (*Cefmo-2*) is both necessary and sufficient to extend health and longevity following hypoxic and metabolic stress^9^. *Cefmo-2* is primarily expressed in the intestine of *C. elegans*, and the FMO family is highly conserved across taxa^11^. *Cefmo-2* is necessary to preserve health during environmental stress and aging in worms. The function, structure, and homeostatic pathways of the nematode intestine are largely conserved, making it a useful model organism to investigate intestinal integrity and stress.

While the xenobiotic roles of FMOs have been well characterized, the endogenous roles of mammalian FMOs are still unclear^12,13^. The mammalian homolog of *Cefmo-2*, Fmo5, is primarily expressed in epithelial cells of the small intestine and colon, as well as in hepatocytes^14,15^.

Fmo5 was recently reported as a regulator of metabolism and nutrient uptake using a constitutive KO mouse model^16^. However, little is known about the functional or cell autonomous role of FMO5 in the intestine. We developed an intestine-specific, tamoxifen-inducible Fmo5 KO mouse to interrogate FMO function in the mammalian intestine. Here, through the establishment of this mouse model, we discovered that Fmo5 plays a critical role in maintaining intestinal homeostasis. We show that disrupting Fmo5 in the adult intestine leads to aberrant GC localization and migration following 14 days of KO. We also find that intestinal Fmo5 is necessary to maintain mucosal barrier thickness in mice, in a sex-dependent manner. We rescue these impairments with an ER stress chaperone, supporting a role for Fmo5 in ER homeostasis. We propose a mechanistic role for Fmo5 in potentiating mucin processing within the ER that is imperative for mucosal barrier homeostasis in mice.

## RESULTS

### *fmo-2* mediates intestinal integrity in *C. elegans*

Based on previous work showing that *Cefmo-2* is necessary and sufficient to improve stress resistance and longevity and that it is expressed primarily in the intestine^9,11^, we were interested whether *fmo-2* affects worm intestinal integrity. To assess intestinal permeability in *C. elegans* we modified an assay commonly utilized in *Drosophila*, the Smurf assay^17^. This technique utilizes a non-absorbable dye that does not cross the intestinal barrier in healthy organisms.

Following intestinal injury in worms, the increased permeability leads to systemic detection of the fluorescent dye (**Fig. 1A**). In wildtype worms, the intestinal permeability of *C. elegans* increases as the worm ages (**Fig. 1B**). This finding is consistent with previous work in aged *C. elegans*, *D. melanogaster*, and mammals showing a gradual decline in intestinal integrity with age. To test whether *Cefmo-2* plays a role in maintaining intestinal integrity, we used the Smurf assay with *fmo-2* knockout (KO) and *fmo-2* overexpressing (OE) worms. We found that worms without *fmo-2* display an increase in intestinal permeability by day 6 of adulthood (**Fig. 1C**). By day 10 of adulthood, wildtype worms showed a similar increase in permeability as *fmo-2* KO worms, whereas *fmo-2* OE worms have less permeability than wildtype worms (**Fig. 1C**). This result suggests that FMO-2 is critical in maintaining intestinal integrity during aging.

**Figure 1.**
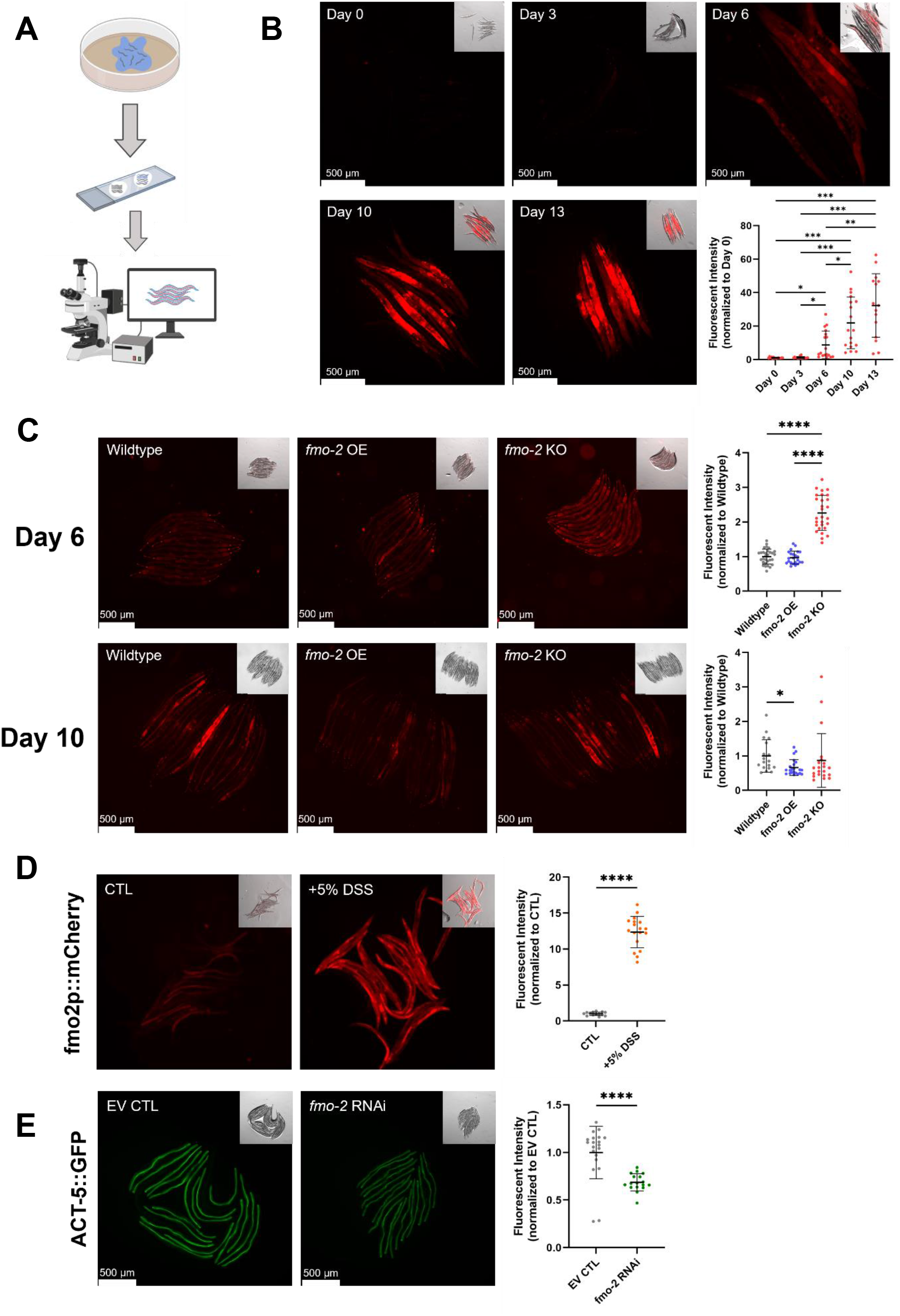
*fmo-2* mediates intestinal integrity and structure in *C. elegans*. (**A**) Diagram of the Smurf assay. (**B**) Fluorescent images of wildtype (N2) worms fed Smurf -dyed (FD&C) food (*E. coli* OP50) at days 0, 3, 6, 10, and 13 of adulthood, and quantification of individual worm fluorescent intensity for each day (n = 15-18 worms/condition). (**C**) Smurf assay of wildtype, *fmo-2* OE, and *fmo-2* KO worms at days 6 (n = 10-16 worms/condition) and 10 of adulthood (n = 18-22 worms/condition), and quantification of individual worm fluorescent intensity for each day. (**D**) Fluorescent images of *fmo-2p::mCherry* worms on day 2 of adulthood following 20 hours of exposure to 5% DSS or M9 control, and quantification of individual worm fluorescent intensity (n = 14-18 worms/condition). (**E**) Fluorescent images of ACT-5::GFP worms on Empty Vector (EV) CTL or *fmo-2* RNAi on day 1 of adulthood, and quantification of individual worm fluorescent intensity (n = 17-20 worms/condition). Lines superimposed onto quantification graphs display mean +/- standard deviation (SD) normalized to Day 0, Wildtype, CTL, or EV CTL, respectively. Scale bars = 500μm. Statistical significance for **B** and **C** was calculated using 1-way ANOVA with Tukey correction for multiple comparisons. Significance for **D** and **E** was calculated by unpaired t-test, two-tailed, with Welch’s correction. * p < 0.05, ** p < 0.01, *** p < 0.001, ****p < 0.0001.

To test whether FMO-2 is also involved in responding to intestinal damage, we utilized dextran sodium sulfate (DSS) to induce intestinal injury^18^. Employing a *fmo-2* transcriptional reporter worm strain, we find that *fmo-2* is significantly induced in worms following 20 hours of exposure to 5% DSS (**Fig. 1D**). Combined with previously published results from our lab and others, where *fmo-2* is induced with exposure to other environmental stressors (dietary restriction, hypoxia, pathogens), this finding is consistent with a model of FMO-2 playing a role in broad environmental stress response^9,19^. To assess the importance of *fmo-2* in maintaining the physical integrity of intestinal cells in *C. elegans*, we used an ACT-5 translational reporter worm strain and RNA interference (RNAi) to knockdown *fmo-2*. ACT-5 is the singular actin protein in the intestinal cells of *C. elegans*, where it makes up the structure of microvilli in the intestinal lumen^20^. Knocking out ACT-5 in worms is embryonically lethal, and the amount of ACT-5 is directly related to the integrity of intestinal cells where lower levels of ACT-5 lead to increased permeability^20^. We find significantly decreased levels of ACT-5 protein in worms when *fmo-2* is knocked down with RNAi (**Fig. 1E**), suggesting *fmo-2* is necessary for maintaining intestinal integrity in the basal state.

### Fmo5 plays a sex-dependent role in maintaining intestinal crypt architecture

There are five FMOs in *C. elegans* and mammals. The phylogenic analysis of active site sequences of FMO proteins suggests that all *C. elegans* FMOs are most closely related to mammalian Fmo5^21^. Fmo5 in mammals is primarily found in the liver and the epithelium of the small intestine and colon (**Fig. 2A**)^15^. To explore the conserved role of *Cefmo-2*/Fmo5 we generated a Fmo5 floxed mouse line (Fmo5^F/F^). Building on our finding that *fmo-2* is necessary for a healthy intestine in *C. elegans*, we crossed the Fmo5 floxed mouse line with an intestinal epithelium-specific, tamoxifen-inducible VillinER^T2^ cre line (Fmo5^IntKO^) (**Fig. 2B**). Gene expression and Western analysis 2 weeks after FMO5 disruption revealed significantly less Fmo5 expression in Fmo5^IntKO^ mice when compared to Fmo5^F/F^ mice (**Fig. 2C, D**).

**Figure 2.**
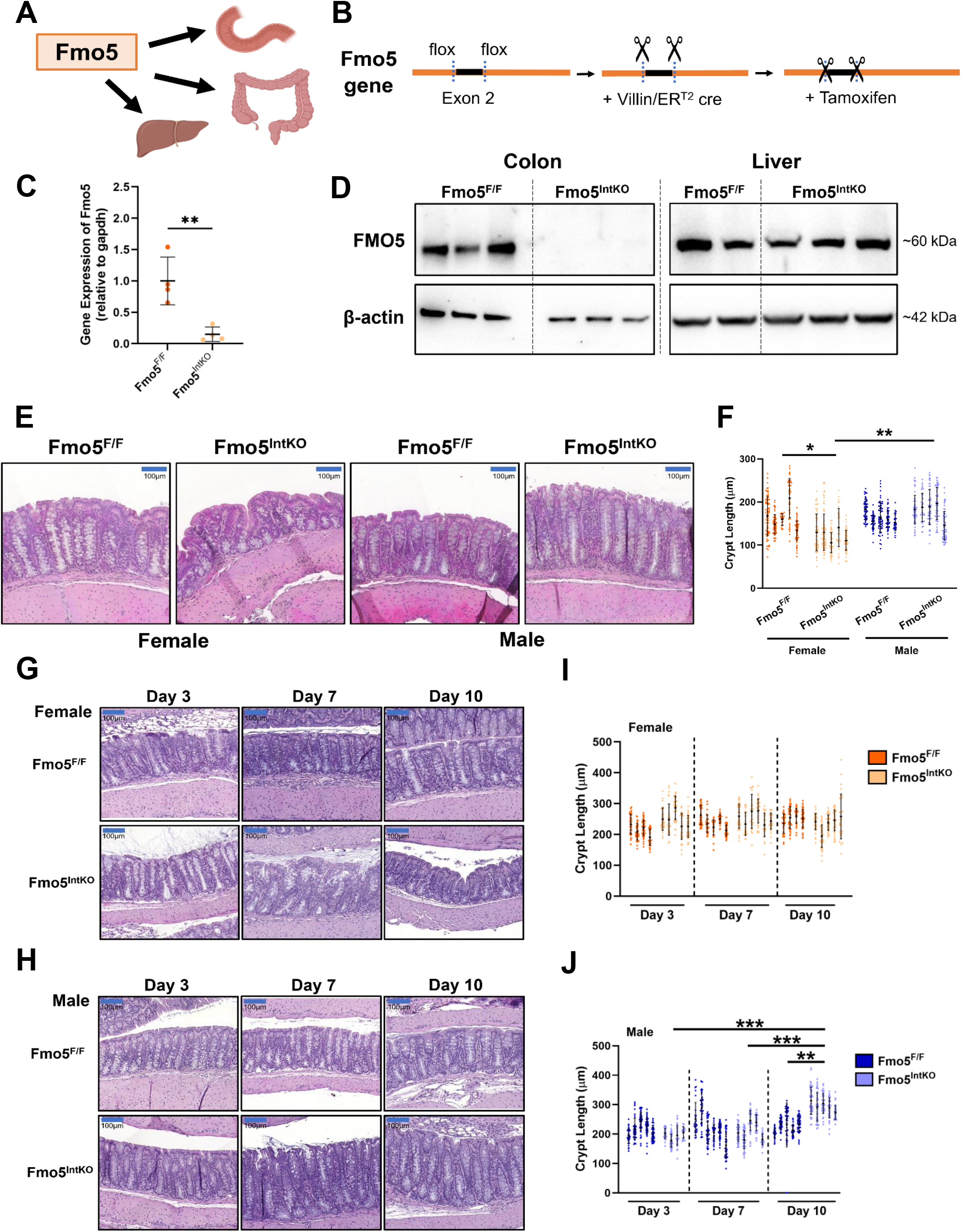
Fmo5 maintains crypt architecture in female mice. (**A**) Graphic of the organs where Fmo5 is expressed in mammals. (**B**) Diagram showing the design of the VillinER^T2^ cre^+^; Fmo5^flox/flox^ mouse line. (**C**) Relative gene expression of Fmo5 in colonic epithelium in Fmo5^F/F^ and Fmo5^IntKO^ mice at 14 days after tamoxifen treatment (n = 4-5 mice/group). Lines superimposed onto plot display mean +/- SD. Statistical significance between groups was calculated using an unpaired t-test, two-tailed. (**D**) Western blot of FMO5 and β-actin proteins in colon epithelium and liver of Fmo5^F/F^ and Fmo5^IntKO^ mice at 14 days of KO. (**E**) Representative H&E-stained images of the proximal colon of female and male Fmo5^F/F^ and Fmo5^IntKO^ mice after 14 days of KO. Scale bar = 100μm, 20x magnification. (**F**) Crypt length quantification (μm) of female and male Fmo5^F/F^ and Fmo5^IntKO^ mice proximal colon represented in **E**. Each bar represents a single mouse where 40-50 crypts were measured. Error bars are displayed as mean +/- SD of measurements from each mouse (n = 5 mice/group). (**G, H**) Representative H&E-stained images of the proximal colon from female (**G**) and male (**H**) Fmo5^F/F^ and Fmo5^IntKO^ mice at days 3-, 7-, and 10- of KO. Scale bar = 100μm, 20x magnification. (**I**, **J**) Crypt length quantification (μm) of female (**I**) and male (**J**) mice represented in **G** and **H**. Each cluster of samples on plots represents a single mouse where 30-40 crypts were measured (n = 4-6 mice/group), with error bars displaying mean +/- SD of measurements from each mouse. A nested 1-way ANOVA with multiple comparisons and Tukey post-hoc correction was performed for data in **F**, **I**, and **J** to determine statistical significance. Significant differences between groups are represented by: * p < 0.05, ** p < 0.01, *** p < 0.001.

Histological analysis of the colonic epithelium in these mice revealed significantly shorter crypts and a dysregulation in the uniformity of crypt structure at 14 days of knockout in female Fmo5^IntKO^ mice, as compared to their Fmo5^F/F^ littermates. In male Fmo5^IntKO^ mice we found elongated crypts compared to male Fmo5^F/F^ mice. This phenotype occurred despite no differences in colonic Fmo5 expression between female and male wildtype mice (**Supplemental Fig. 1A**). Sex differences have been noted in several gastrointestinal diseases in humans^22^.

To determine if Fmo5 plays a role in maintaining crypt architecture and/or if Fmo5 is necessary for epithelial cell turnover in the colon, we performed H&E staining in Fmo5^IntKO^ and Fmo5^F/F^ mice at 3-, 7-, and 10- days following tamoxifen treatment. After 3 days of knockout, we did not see distinct differences in epithelial architecture between female Fmo5^IntKO^ and Fmo5^F/F^ mice. However, 7 days following the disruption of Fmo5 in the intestine there was a loss of crypt uniformity in female Fmo5^IntKO^ mice (**Fig 2G**). Crypts were stacked on top of one another causing the epithelium to appear increasingly disorganized (**Fig 2G**). At day 10 of KO, the uniformity of crypt architecture remained lost in these mice.

Similar to what we observed in females, there were no differences between male Fmo5^F/F^ and Fmo5^IntKO^ mice at day 3 and 7 of KO. At 10 days post knockout, male Fmo5^IntKO^ mice had significantly longer crypts than both their wildtype littermates and Fmo5^IntKO^ mice at days 3 and 7 (**Fig. 2H, J**). The absence of significant epithelial changes in both female and male Fmo5^IntKO^ mice after 3 days indicates that Fmo5 is not essential for acute cell survival but rather affects the subsequent turnover and development of the epithelium.

### Female Fmo5^IntKO^ mice have impaired goblet cell localization and maturation

The intestinal epithelium is made up of a heterogenous population of epithelial cells including enteroendocrine, absorptive, and secretory cell types each performing distinct functions to maintain homeostasis^23,24^. We found that the expression of the GC gene Muc2 was significantly increased in female Fmo5^IntKO^ mice (**Fig. 3A**). Also, Tff3 was increased but not statistically significant, but this was not observed in male Fmo5^IntKO^ mice (**Supplemental Fig. 1B**). We did not find significant differences in absorptive enterocyte (Slc2a5 and Alpi), Paneth (Mmp7 and Lys1), tuft (Pou2f3), enteroendocrine (Neurog3 and ChgA), or stem cell (Lgr5 and Bmp4) markers (**Supplemental Fig. 1B**). Immunofluorescence (IF) staining of Muc2 (green) revealed altered localization of GCs in female Fmo5^IntKO^ mice, as GCs are clustered at the bottom of the crypt compared to the Fmo5^F/F^ mice where GCs are evenly distributed throughout the crypt (**Fig. 3B**).

**Figure 3.**
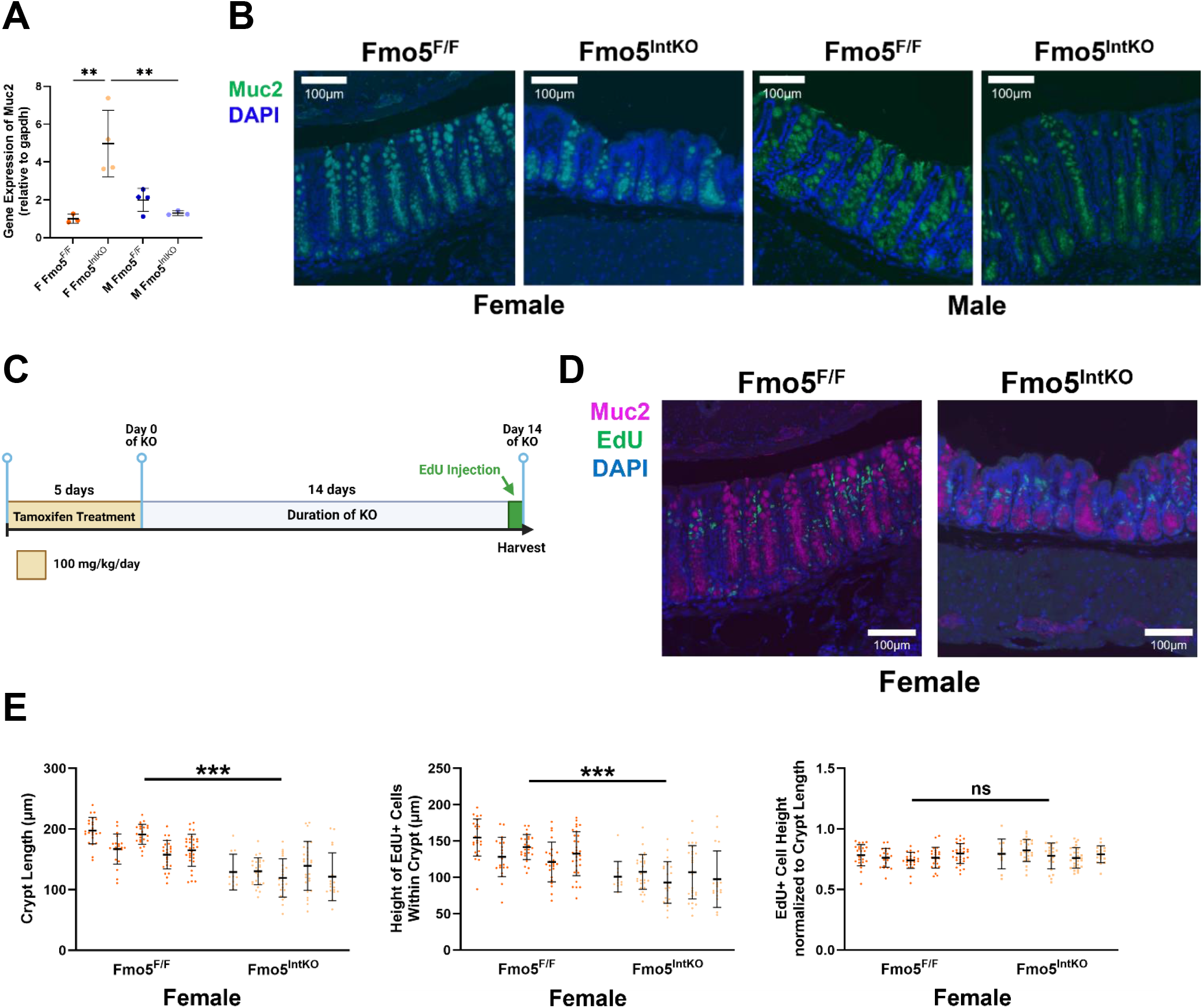
Loss of Fmo5 impairs goblet cell migration. (**A**) Relative gene expression of Muc2 in colonic epithelium of female (F) and male (M) Fmo5^F/F^ and Fmo5^IntKO^ mice 14 days after tamoxifen treatment (n = 3-4 mice/group). Lines superimposed onto plot display mean +/- SD. (**B**) Representative images of immunofluorescence staining of Muc2 (green) and DAPI (blue) in female and male Fmo5^F/F^ and Fmo5^IntKO^ proximal colons at day 14 of KO. Scale bar = 100μm, 20x magnification. (**C**) Experimental design of tamoxifen treatment and EdU labeling injection. (**D**) Representative images of Muc2 (magenta), EdU (green), and DAPI (blue) stained colons collected 24 hours after EdU labeling. Scale bar = 100μm, 20x magnification. (**E**) Quantification of crypt length (μm) and the respective height of EdU+ cells within that crypt 24 hours after EdU labeling (day 14 of KO). Quantified EdU+ cell migration height (μm) normalized by each specific crypts’ length (μm). Each cluster of samples represents a single mouse where 10-25 crypts/mouse were measured (n = 5 female mice/group), with error bars displaying mean +/- SD. A nested 1-way ANOVA with multiple comparisons and Tukey post-hoc correction was performed for data in **A**. Unpaired t-tests were used to calculate statistical significance between groups in **E**. ** p < 0.01, *** p < 0.001.

To determine if female Fmo5^IntKO^ mice have an impairment in overall epithelial cell migration, we performed a pulse chase experiment in Fmo5^F/F^ and Fmo5^IntKO^ mice with EdU, for 24 hours before collecting the colon for histological analysis (**Fig. 3C**). We then performed IF staining of Muc2 (magenta) and EdU (green) (**Fig. 3D**) to visualize the location of cells that were actively replicating when EdU was injected 24 hours prior. We measured the distance up the crypt that the EdU+ cells had migrated within 24 hours and normalized that distance to individual crypt length (**Fig. 3E**), finding no difference in the migration distance of EdU+ cells between female Fmo5^F/F^ and Fmo5^IntKO^ mice at 24 hours. This supports the conclusion that overall epithelial cell migration is not altered in female Fmo5^IntKO^ mice, but that there is a migration impairment specific to GCs. Goblet cells mature as they migrate up the crypt towards the lumen of the intestine, which is required to properly secrete mucin in response to systemic cues^25,26^. In support of this, we found no difference in absorptive or secretory progenitor cell expression (Hes1, Atoh1) in our mice, suggesting the lack of goblet cells in the upper crypt is not due to a decrease in secretory cell lineage differentiation, but rather an impairment in GC maturation (**Supplemental Fig. 1C**). These results, in conjunction with the spatial impairments of GCs within the crypts observed in female Fmo5^IntKO^ mice, support a role for Fmo5 in GC development and maturation.

### Fmo5 is necessary for mucus production and barrier thickness in female mice

Muc2 protein undergoes complex folding and N-glycosylation within the ER^27^ before moving to the Golgi apparatus to be O-glycosylated^28^. Following post-translational modification, mucin is stored in secretory vesicles in the cup-shaped region of the cell to await release into the crypt or luminal space. Given that GC localization and maturity is impaired in female Fmo5^IntKO^ mice, we next wanted to test the functional capacity of GCs to produce mucus. AB/PAS staining of mucus within the crypts of Fmo5^F/F^ and Fmo5^IntKO^ mice revealed a decrease in overall mucus production from GCs within female Fmo5^IntKO^ crypts. This suggests that the spatial and migration impairments of GCs in these mice are sufficient to negatively impact their capacity to produce mucus (**Fig. 4A**). We observed a slight reduction in mucus within the crypts of male Fmo5^IntKO^ mice, although not to the severity observed in females. We find that female Fmo5^IntKO^ mice had similar levels of mucus within the crypt as their wildtype littermates at day 3 post-tamoxifen treatment. By day 7, female KO mice began to show dysregulated crypt organization along with noticeably less mucus within the epithelium (**Fig. 4B**). At day 10 of KO, female Fmo5^IntKO^ mice displayed shorter crypts, similar to what we saw at day 14 of KO. This is coupled with a lack of mucus in the upper half of crypts, whereas in Fmo5^F/F^ mice mucus-filled GCs spread from the apical to basolateral borders of the crypts. There were no differences in mucus production in male Fmo5^IntKO^ mice at day 3 of KO (**Fig. 4C**). At day 7 post-tamoxifen, there was a slight reduction in mucus present within the crypts of male Fmo5^IntKO^ mice when compared to Fmo5^F/F^. This reduction was also present at day 10 and 14 of KO, where there was mild loss of mucus sporadically among crypts (**Fig. 4C**).

**Figure 4.**
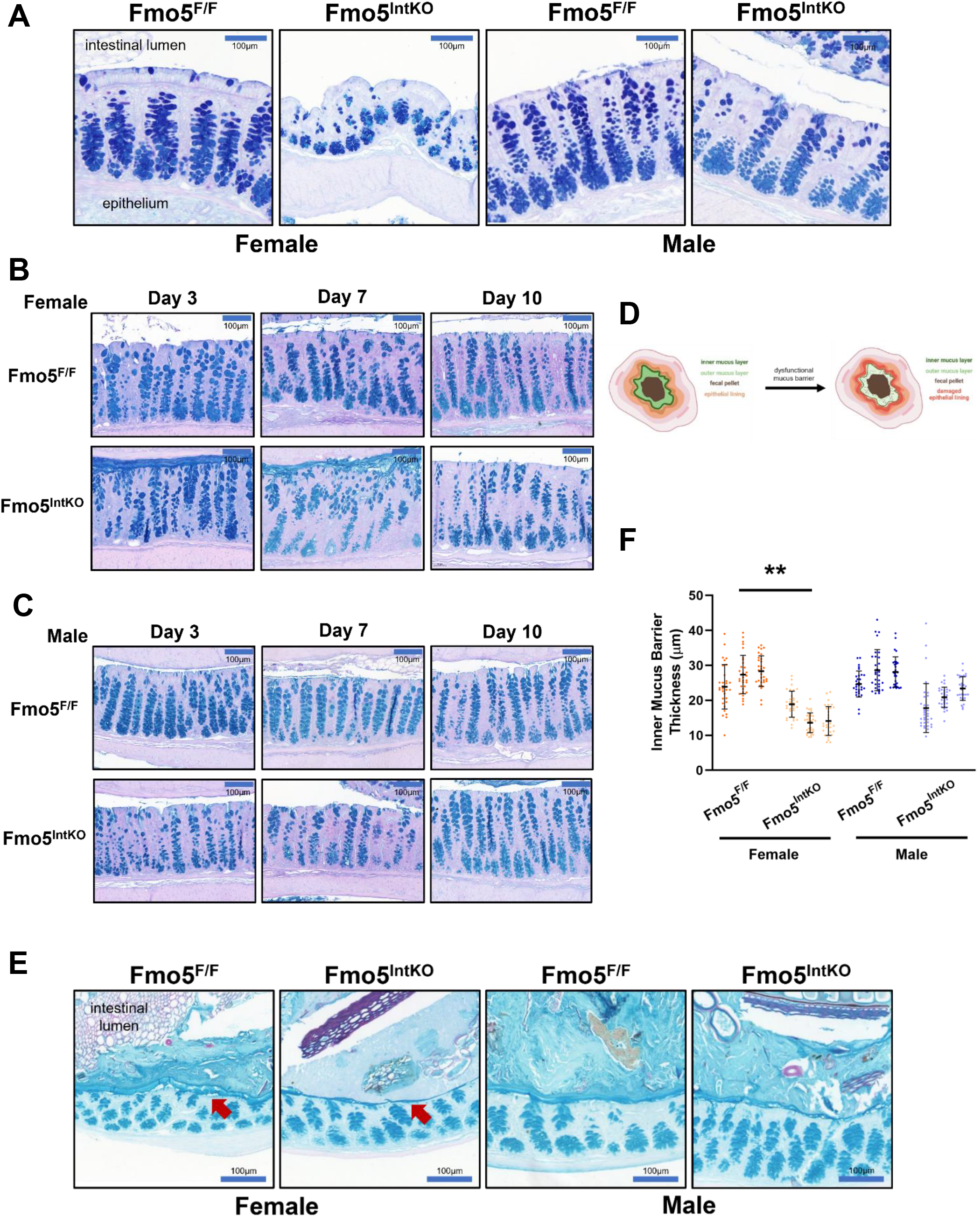
Loss of Fmo5 decreases mucus barrier thickness in female mice. Representative images of AB/PAS-stained proximal colon samples from female and male Fmo5^F/F^ and Fmo5^IntKO^ mice at (**A**) day 14 of KO, (**B**) females at days 3-, 7-, and 10- of KO, and (**C**) males at days 3-,7-, and 10- of KO. All images were taken at 20x magnification. Scale bar = 100μm. (**D**) Schematic of cross-sectional view of the colon with healthy vs. dysfunctional mucus barrier. (**E**) Representative images of colon cross-sections stained with AB/PAS and processed using Carnoy’s fixative to preserve luminal mucus barrier in female and male mice 14 days following tamoxifen treatment, 20x magnification. Scale bar = 100μm. Red arrows point to the inner mucus barrier. (**F**) Quantified inner mucus barrier thickness (μm) of images represented in **E**. ∼30 measurements of inner mucus barrier thickness were made for each mouse (n = 3 mice/group), where each cluster of samples represents data from one mouse. Error bars display mean +/- SD for each mouse. Statistical significance was calculated using nested 1-way ANOVA with multiple comparisons and Tukey post-hoc correction. * p < 0.05, ** p < 0.01.

Histological preparation of the colon removes the luminal mucus barrier but preserves intracellular mucus in GCs. Interestingly, we observed that all female Fmo5^IntKO^ mice at day 3 of KO have a layer of mucus adhered to the epithelium that was not disturbed during processing. Additionally, mucus was present in 4/5 female Fmo5^IntKO^ mice at day 7 of KO, which was not observed in any Fmo5^F/F^ or male Fmo5^IntKO^ mice. This observation suggests that there are differences in mucus composition in female Fmo5^IntKO^ that prevent the mucus layer from being washed away during standard processing. Using Carnoy’s fixative solution that preserves the mucus barrier (**Fig. 4D**), we found a significantly thinner inner mucus layer in female Fmo5^IntKO^ mice at day 14 of KO when compared to female Fmo5^F/F^ mice (**Fig. 4E, F**). Overall, these results demonstrate both impaired mucus production within crypts and the necessity of Fmo5 to maintain mucus barrier thickness in female mice.

### Mucus impairments in Fmo5^IntKO^ mice are not likely caused by dysbiosis

To determine if the damaged epithelium and dysregulated mucus barrier in female Fmo5^IntKO^ mice is due to dysbiosis, we utilized 16s rRNA sequencing of feces from female and male Fmo5^F/F^ and Fmo5^IntKO^ mice. We found no significant changes in the microbial populations of Fmo5^IntKO^ animals compared to their littermate controls after 13 days of KO (**Fig. 5A**), indicating that dysbiosis is not likely the primary cause of the intestinal phenotype. Moreover, Fmo5^F/F^ and Fmo5^IntKO^ mice treated with broad-spectrum antibiotics (**Fig. 5B**) did not rescue crypt damage or GC alterations in female Fmo5^IntKO^ mice (**Fig. 5C, D, E**). When comparing crypt structure in Fmo5^F/F^ and Fmo5^IntKO^ mice, both wildtype groups and male Fmo5^IntKO^ mice experience crypt shortening with antibiotics. In contrast, female Fmo5^IntKO^ crypts do not shorten in response to antibiotics.

**Figure 5.**
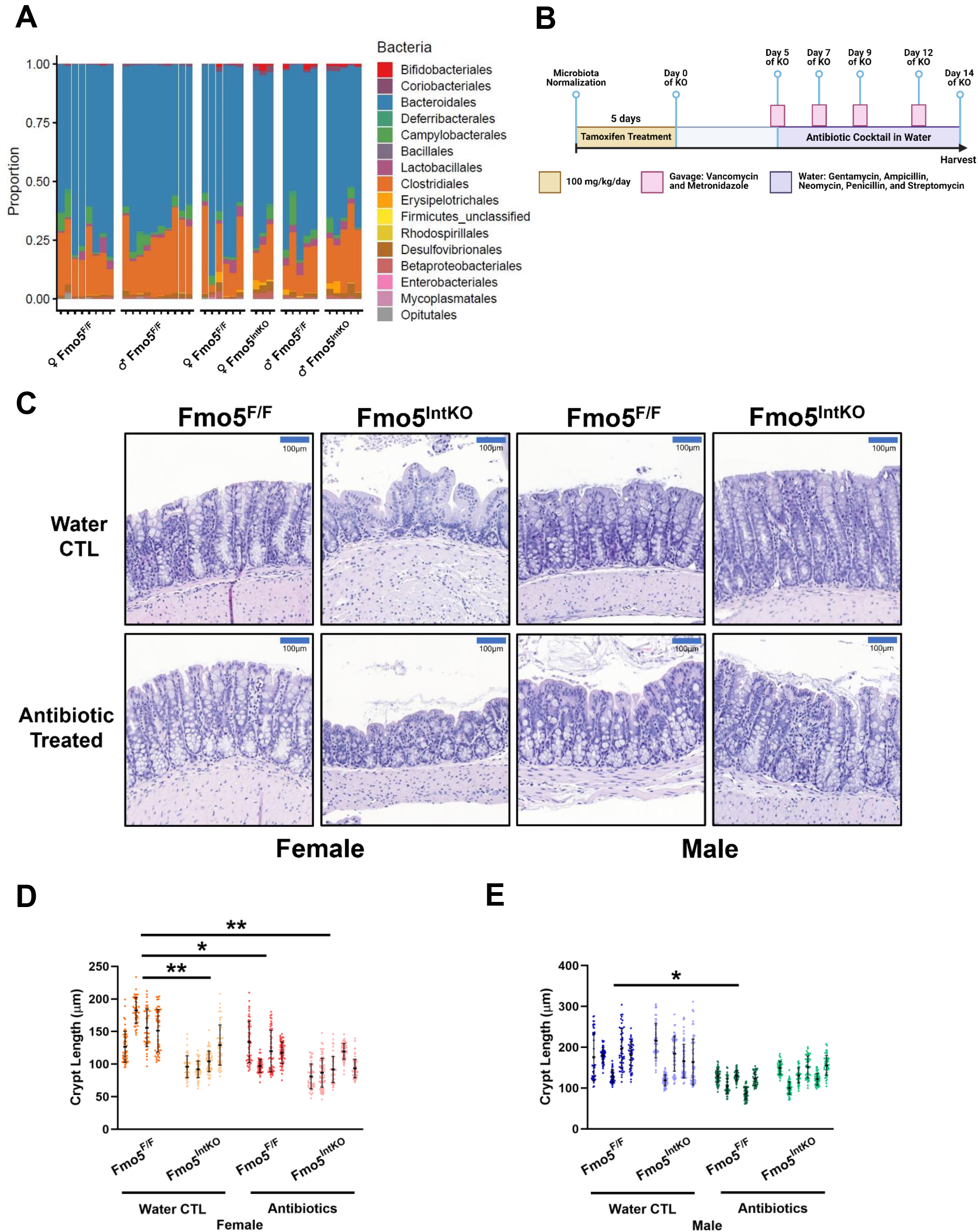
No evidence of dysbiosis in female Fmo5^IntKO^ mice. (**A**) Graph displaying bacterial 16s rRNA sequencing of fecal pellets from female and male Fmo5^F/F^ mice prior to tamoxifen treatment, and female and male Fmo5^F/F^ and Fmo5^IntKO^ mice after 13 days of KO (n = 3-6 mice/group). ♀=female, ♂=male. (**B**) Experimental design and timeline of tamoxifen and antibiotic treatment. (**C**) Representative images of H&E-stained proximal colons from female and male Fmo5^F/F^ and Fmo5^IntKO^ mice after 14 days of KO. Antibiotic-treated mice were given a mixture of Gentamycin, Ampicillin, Neomycin, Penicillin, and Streptomycin in drinking water beginning at 5 days post-tamoxifen treatment. Mice were also gavaged with Vancomycin and Metronidazole (Antibiotic Treated) or PBS (Water CTL) every other day from 5-14 days of KO. 20x magnification, Scale bar = 100μm (**D, E**) Quantification of crypt length (μm) from (**D**) female and (**E**) male mice described in **C**. ∼45-60 crypts were measured from the proximal colon of each mouse. Each cluster of data represents measurements from one mouse (n = 4-6 mice/group), and error bars reflect mean +/- SD for that mouse. Statistical significance was determined using a nested 1-way ANOVA with Tukey-corrected multiple comparisons. * p < 0.05, ** p < 0.01.

### The ER chaperone TUDCA rescues crypt dysregulation and mucus barrier impairment in female Fmo5^IntKO^ mice

The capacity of the ER to properly fold the large Muc2 protein during mucin synthesis is imperative for maintaining a healthy intestinal mucus barrier. There were no significant differences in the expression of key players in UPR^ER^ stress response (BIP, CHOP, IRE1α, PERK, ATF6, ATF4) (**Supplemental Fig. 2A**) or in BIP or PERK protein levels (**Supplemental Fig. 2B, C**). In conjunction with this, we see no differences in Spliced- or Total XBP1 expression (**Supplemental Fig. 2D**), suggesting the lack of increased canonical UPR^ER^ activation in female Fmo5^IntKO^ mice. Considering how imperative it is for GCs to effectively complete the many steps involved in producing and secreting mucus, the ER of GCs has evolved to include an additional UPR^ER^ arm composed of the mucus-specific ER stress sensor IRE1β (inositol-requiring enzyme 1 beta) and it’s chaperone AGR2 (anterior gradient protein 2 homolog)^29,30^. AGR2 levels were significantly higher in Fmo5^IntKO^ mice when compared to Fmo5^F/F^ mice (**Fig. 6A, B**). This was coupled with significantly increased protein levels of IRE1β in female Fmo5^IntKO^ mice (**Fig. 6A, B**), indicating activation of the mucus-specific ER stress response pathway. To explore if mucus-specific ER stress is contributing to the crypt and mucus barrier defects that we see in female Fmo5^IntKO^ mice, we treated Fmo5^F/F^ and Fmo5^IntKO^ mice with the bile acid chemical ER chaperone Tauroursodeoxycholic acid (TUDCA). TUDCA or PBS (control) was given 7 days prior to and during tamoxifen treatment, in addition to 15 days following KO (**Fig. 6C**). H&E staining to assess crypt architecture revealed a complete rescue of crypt length in female Fmo5^IntKO^ mice with TUDCA treatment, compared to Fmo5^IntKO^ mice given PBS (**Fig. 6D, E**).

**Figure 6.**
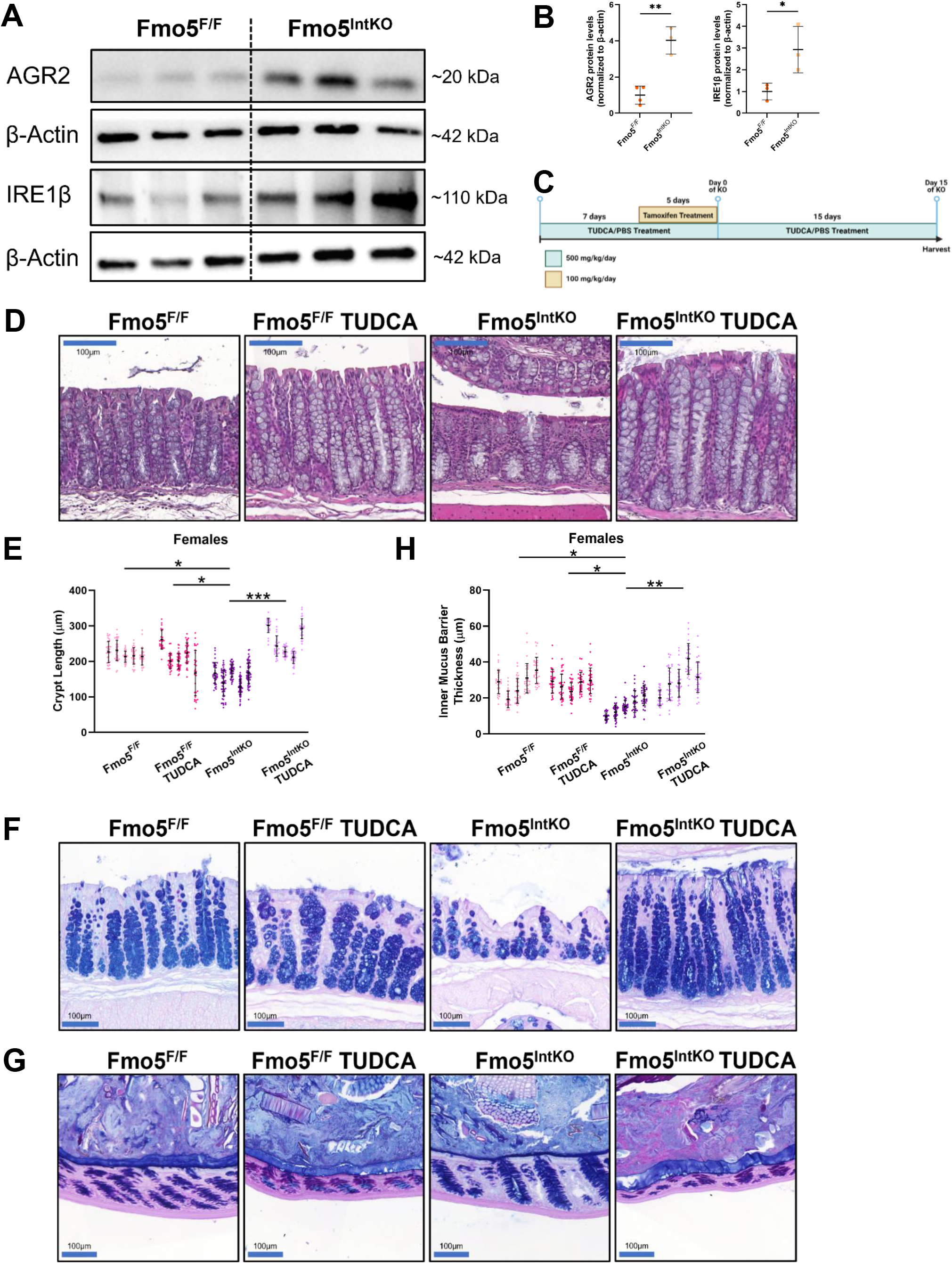
TUDCA treatment restores crypt length and mucus barrier thickness in female Fmo5^IntKO^ mice. (A) Representative Western blots of Agr2 and IRE1β proteins with β-actin controls from female Fmo5^F/F^ and Fmo5^IntKO^ mice at 14 days of KO. (**B**) Quantification of Agr2 and IRE1β protein levels described in **A**, normalized to their respective β-actin levels (n = 3-4 mice/group). Lines superimposed on plot display mean +/- SD. (**C**) Experimental design and timeline of TUDCA experiment. (**D**) Representative H&E-stained proximal colon from female Fmo5^F/F^ and Fmo5^IntKO^ mice treated with TUDCA or PBS (CTL). 20x magnification, Scale bar = 100μm. (**E**) Crypt length (μm) quantification of samples represented in **D**. ∼30 crypts were measured from each mouse with clusters of samples representing measurements from one mouse (n = 5 mice/group). Error bars display mean +/- SD for each mouse. (**F**) AB/PAS-stained images of female mice +/- TUDCA treatment. 20x magnification, Scale bar = 100μm. (**G**) Representative images of AB/PAS-stained colon cross-sections from TUDCA experiment that were processed using Carnoy’s fixative. 20x magnification, Scale bar = 100μm. (**H**) Quantification of inner mucus barrier thickness (μm) represented in **G**. ∼30 measurements of inner mucus barrier thickness were made for each mouse. Each cluster of samples displays data from one mouse with error bars showing mean +/- SD for that mouse (n = 5 mice/group). Statistical significance was calculated by unpaired t-test (**B**) and nested 1-way ANOVA with Tukey correction for multiple comparisons (**E, H**). Significant differences between groups are represented by: * p < 0.05, ** p < 0.01, *** p < 0.001.

Mucus production from GCs within the crypt was also restored with TUDCA treatment, where female Fmo5^IntKO^ mice given TUDCA have visually comparable, if not increased, amounts of intracellular mucus present (**Fig. 6F**). Additionally, we found that TUDCA treatment in Fmo5^IntKO^ mice completely restored inner mucus barrier thickness when compared to Fmo5^IntKO^ mice without TUDCA (**Fig. 6G, H**). The ability of an ER chaperone to improve crypt and mucus defects in mice without Fmo5 suggests that the underlying cause of these defects involves protein folding and/or the ER stress response.

## DISCUSSION

Together, our results support a model in which FMOs play a conserved, integral role in maintaining intestinal homeostasis. In *C. elegans,* we show that *fmo-2* is necessary to maintain structural integrity and barrier function in the intestine (**Fig. 1**). Building upon this, to define the conserved role of FMOs in maintaining intestinal homeostasis, we generated a tamoxifen-inducible, intestine-specific mouse model of mammalian Fmo5. Characterization of this mouse line revealed a sex-dependent role for Fmo5 in the maintenance of epithelial architecture in the colon (**Fig. 2E**). Our results also demonstrate the necessity of Fmo5 for GC localization and Muc2 expression, leading to thinning of the mucosal barrier with loss of Fmo5 in mice (**Fig. 4E, F**). High demand on the GC ER to process mucins and evidence of mucus-specific ER stress in Fmo5^IntKO^ animals led us to hypothesize that Fmo5 is essential to maintain ER homeostasis during mucus production. This hypothesis is supported by the rescue of crypt architecture and mucus barrier thickness with the addition of the bile acid ER chaperone, TUDCA (**Fig. 6D, G**). Our results 1) support a new role for FMO5 in maintaining intestinal ER homeostasis, and 2) introduce a new mouse model of intestinal dysfunction. Overall, this work highlights the importance of Fmo5 in maintaining a healthy intestine in mice.

Our results show the highly conserved enzyme, FMO5, when disrupted in mice, exhibits many characteristics of chronic GI disease, but only in female mice. This is a striking sex difference that parallels that observed in humans, despite no clear linkage between Fmo5 and GI disease pathogenesis. The mechanism underlying sexual dimorphism in GI disease is largely unclear, and this model provides a unique opportunity within the field to dissect the cellular mechanisms. Both the prevalence and progression of IBD differs in humans based on sex, where women are more likely to develop Crohn’s Disease (CD) and men are more susceptible to ulcerative colitis (UC)^22^. One study found that post age 45, females are up to 32% less likely to be diagnosed with UC when compared to males^31^. This is coupled with another report showing that Fmo5 expression was down-regulated in a UC dataset^32^. Interestingly, we found no differences in Fmo5 expression levels in the intestinal epithelium between female and male mice at 1, 3, or 6 months of age (**Supplemental Fig. 1A**). This finding does not support the hypothesis that female mice are more susceptible to Fmo5 loss due to higher intestinal Fmo5 levels. We speculate that Fmo5 may interact with sex hormones to provide protection from intestinal dysfunction in female mice.

Maintaining the physical integrity of the intestine by epithelial cells is critical for preserving intestinal homeostasis and function. Morphological assessment of crypt architecture is a useful tool when diagnosing GI disease in a clinical setting, and its relevance to disease spans into mammalian models^33,34^. Studies show that shortening of crypts in the colon (crypt atrophy), crypt distortion, and crypt loss are all present in mouse models of IBD and these alterations have a direct impact on epithelial health^35^. Additionally, crypt shortening and disarray along with crypt branching and mucin depletion are diagnostic features of UC in humans^36–39^. These characteristics are observed in the colonic epithelium of female Fmo5^IntKO^ mice, leading us to hypothesize that Fmo5 plays a protective role in GI disease onset. The severity of crypt shortening that we report (**Fig. 2E**) is comparable to mice following acute colitis^35^. Crypt lengthening in male Fmo5^IntKO^ mice could be attributed to crypt epithelial hyperplasia or crypts elongating to increase cell number and surface area to maintain epithelial function in Fmo5^IntKO^ mice (**Fig. 2F**)^40,41^. We also found that the loss of structural homeostasis in the colon of female Fmo5^IntKO^ mice compounds with each epithelial turnover, resulting in dramatic crypt distortion by 14 days post-KO (**Fig. 2G**). From this, we conclude that Fmo5 is necessary for the formation of a healthy epithelium in the colon. The absence of further crypt shortening in female Fmo5^IntKO^ mice with antibiotic treatment could be due to the loss of beneficial bacteria in the gut. We speculate that female Fmo5^IntKO^ mice are severely compromised, and that any greater loss of crypt architecture will be detrimental to survival. Despite the lack of evidence linking the Fmo5^IntKO^ phenotype to dysbiosis, it is possible that female Fmo5^IntKO^ mice have different interactions with homeostatic gut bacteria. This possibility is supported by published work on a Fmo5 constitutive knockout animal model, where male mice were suggested to respond differently to microbiota^42^. Additional investigation is needed to determine if female mice without Fmo5 are hypersusceptible to intestinal cues or intestine-specific stress.

Studies show that altered epithelial architecture in mice can result from defects in cell differentiation and migration, where cells are unable to terminally differentiate or effectively migrate up the crypt toward the lumen of the intestine^25^. We find that GCs in female Fmo5^IntKO^ mice are clustered toward the bottom of the crypt rather than being spread uniformly throughout (**Fig. 3B**). This is intriguing, as mice lacking the primary colonic GC mucin (Muc2) exhibit altered crypt morphology and have impairments in GC migration up the crypt. These data are consistent with Scott *et al.,* who observed decreased GC number in constitutive, whole body male Fmo5^-/-^ mice^42^. Multiple factors necessary to drive GC maturation and migration have been identified, and several studies link impairments in these with the development of IBD^39,43^. One phenotype specifically attributed to cases of UC in humans, rather than IBD as a whole, is a loss of mucin-producing GCs in the top half of the crypt coupled with depleted mucus barrier^37,44–46^.

Additionally, GC clustering and loss of upper crypt GCs in unchallenged female Fmo5^IntKO^ mice (**Fig. 3B**) is consistent with published models of DSS-induced colitis, where GC dysfunction is attributed to decreased GC maturation^39,43^. The lack of differences in absorptive and secretory progenitor cell expression in our mice (**Supplemental Fig. 1C**) rule out the possibility that impaired mucus production in female Fmo5^IntKO^ mice is caused by decreased secretory cell differentiation. This further supports the hypothesis that maturation is the likely GC defect occurring in female Fmo5^IntKO^ mice. These findings, coupled with an increase in GC gene expression (Muc2, Tff3) that we suspect is compensatory (**Fig. 3A**, **Supplemental Fig. 1B)**, suggest that Fmo5 contributes to GC regulation in female mice.

Maintaining a protective mucus barrier is the culmination of a range of structural, functional, and biochemical processes. Here we show that intestinal Fmo5 is necessary for proper mucus production within colonic crypts and mucus barrier maintenance in female mice (**Fig. 4A, E**).

Previously published studies show that impairments in Muc2 folding, defects in O- and N-glycosylation, and aggregation of the non-glycosylated form of Muc2 in mice each cause hyper- susceptibility to colitis coupled with aberrant GC function^47^. It is likely that one or more of these protein processing impairments occur in Fmo5^IntKO^ females, given that 1) Fmo5 resides on the ER membrane, and 2) we find evidence of mucus-related ER stress. A previous study showed that the BIP/IRE1α arm of the UPR senses a wide variety of ligands that cause ER stress but does not respond to excess or misfolded Muc2 presence^29^. The mucin-sensitive UPR^ER^ arm made up of AGR2 and IRE1β is hypothesized to take on the responsibility for the proper folding of mucins in GCs^29,48^. This may prevent the canonical ER stress response from overwhelming the ER in mucosal epithelial cells. Our findings are consistent with this, as we show increased AGR2 and IRE1β protein levels in female Fmo5^IntKO^ mice (**Fig. 6A, B**), in the absence of typical UPR^ER^ activation (**Supplemental Fig. 2**). Complete rescue of crypt architecture and mucosal barrier impairments with the addition of the ER chaperone TUDCA (**Fig. 6D-H**) implicates protein processing dysfunction in the ER of female Fmo5^IntKO^ mice. This finding leads us to speculate if Fmo5 loss causes an accumulation of Muc2 proteins in the GC ER that are unable to fold correctly, triggering AGR2 and IRE1β activation. While further exploration of this system is necessary to identify the exact molecular mechanism of how Fmo5 plays into ER stress regulation, our results imply that GC and mucus barrier defects in female Fmo5^IntKO^ mice are caused by altered protein folding and/or ER stress. This establishes a role for Fmo5 in maintaining mucosal barrier integrity and intestinal homeostasis in mice through ER stress protection.

Together, this work describes a previously unidentified role for Fmo5 in maintaining intestinal integrity and mucosal barrier homeostasis. There remains a need to identify the cellular mechanism(s) by which acute loss of Fmo5 leads to defects in GC maturation and migration, and if an altered metabolic environment is involved. Additional investigation is also warranted to determine if there are any changes to the biochemical components of mucus produced in mice without FMO5 and how those changes influence mucus barrier integrity and resilience. Future directions for this project include clarifying the cellular mechanism of how intestinal Fmo5 influences GI disease onset, with the overall goal to utilize the resulting data to improve therapeutic treatment for GI diseases.

## MATERIALS AND METHODS

### Worm Strain and Maintenance

Wildtype N2, KAE9 (*fmo-2* OE), VC1668 (*fmo-2* KO), ELT-60 (ACT-5::GFP), and LZR1 (*fmo-2p::MC*) strains of *C. elegans* were age synchronized by timed egg lay prior to use. Strains were grown and maintained on solid nematode growth media (NGM) throughout their lifespan. The nematodes were fed *E. coli* OP50 throughout life except where RNAi (*E. coli* HT115 *fmo-2* and Empty Vector Control) were fed instead. All experiments were performed at 20°C.

### Smurf Assay in *C. elegans*

Wildtype, *fmo-2* KO, and *fmo-2* OE worms were synchronized and fed *E. coli* OP50 bacteria dyed with 0.10% wt/vol. Erioglaucine disodium salt (Sigma Aldrich, FD&C). On each day of analysis, worms were placed on a 3% agarose pad on a glass slide, and 1M sodium azide was used to induce paralysis before imaging using a Leica fluorescent microscope. Fluorescent intensity of each worm was quantified using ImageJ.

### DSS Assay in *C. elegans*

*fmo-2p::mCherry* worms were age synchronized and fed with *E. coli* OP50 until day 1 of adulthood. They were then incubated in M9 wash solution with OP50 +/- 5% dextran sodium sulfate (DSS) for 20 hours at 20°C prior to imaging. Worms were placed on a 3% agarose pad on a glass slide, and 1M sodium azide was used to induce paralysis before imaging using a Leica fluorescent microscope. Fluorescent intensity of each worm was quantified using ImageJ.

### Mouse Line Generation

The Fmo5^F/F^ mouse line was generated in collaboration with the Transgenic Animal Model Core of the University of Michigan’s Biomedical Research Core Facilities. Briefly, purified DNA was microinjected into fertilized eggs obtained by mating (C57BL/6 X SJL) F1 or C57BL/6 female mice with (C57BL/6 X SJL) F1 male mice. Pronuclear microinjection was performed as described^49^. Fmo5^F/F^ were then crossed into the intestine-specific, inducible VillinER^T2^ cre mouse line and backcrossed a minimum of 5 generations prior to using for experiments.

### Mouse Husbandry

All mice used in this study were 8-12 week old VillinER^T2^ cre^+^; Fmo5^flox/flox^ (Fmo5^IntKO^) and Fmo5^flox/flox^ (Fmo5^F/F^) female and male mice in a C57Bl/6 background. Littermate controls were used whenever possible. All mice were housed in the pathogen-free Unit for Laboratory Animal Management at the University of Michigan. Mice were kept under a 12-hour light/12-hour dark cycle and standard chow and water were provided *ad libitum.* All animal studies were conducted in accordance with Association for Assessment and Accreditation of Laboratory Animal Care International guidelines and approved by the University Committee on the Use and Care of Animals at the University of Michigan.

### Tamoxifen Treatment

Mice were given intraperitoneal injection of 100 mg/kg tamoxifen (MP Biomedical) in corn oil once daily for 5 days. Mice were weighed daily during treatment to monitor health and were housed by genotype and sex. The 5^th^ and final day of injection was considered day 0 of Fmo5 knockout for all experiments in this study. Both female and male mice were used for each experiment except for Figure 3D, 3E, and Figure 6, where only female mice were used. Over the course of the study we observed a phenotype that was sex-dependent, and the listed experiments were primarily relevant to the phenotype observed in female mice.

### Histology

To prepare samples for general histological analysis, the colon was removed at necropsy and opened lengthwise. The colon was rinsed in 1x PBS and rolled to create a swiss roll^50^. Samples were incubated in 10% buffered formalin phosphate (Fisher Chemical) overnight at room temperature before being moved to 70% ethanol until further processing. Tissues were dehydrated by paraffin wax embedding and sliced into 4 µm sections^51^.

For mucus barrier histological analysis, upon euthanasia the colon was removed and kept intact. Horizontal cuts were made above and below a fecal pellet midway down the colon and the sample was immediately placed into Carnoy’s fixative (Spectrum) to preserve the mucus barrier^52^. Samples were incubated overnight at room temperature in Carnoy’s solution prior to being moved to 70% ethanol and stored at 4°C until paraffin embedding. Embedding and sectioning was performed using the same method as described above.

### Immunofluorescence

Slides were deparaffinized using xylene and rehydrated using washes of 100%, 95%, and 80% ethanol at room temperature. Samples underwent antigen retrieval in sodium citrate buffer with a pH of 6.0 (Thermo Scientific) at 100°C for 20 minutes. Samples were blocked with goat serum and incubated in Muc2 primary antibody (1:1000, Proteintech) overnight at 4° at a concentration of 1:1000. The following day, slides were rinsed and incubated in secondary antibody (Anti-GFP, Invitrogen) for 1 hour at room temperature (RT) at 1:500 concentration. Following incubation, samples were washed, and cover slips were mounted using Prolong Gold with DAPI (Fisher Chemical).

### EdU Injection and Staining

Mice were given intraperitoneal injection of 25 mg/kg of EdU (5-ethynyl-2’-deoxyuridine, Invitrogen) dissolved in 1x PBS on day 13 of knockout. Exactly 24 hours later, mice were euthanized by C02 inhalation, and intestinal tissue was collected for histological analysis. Samples were deparaffinized using xylene and rehydrated using washes of 100%, 95%, and 75% ethanol at room temperature. Immunofluorescence staining as described above was performed until after the secondary incubation step. Slides were then incubated in 0.5% Triton X-100/PBS for 20 minutes at RT and EdU staining was performed using a Click-iT EdU Kit (Invitrogen) following the included protocol. Briefly, slides were incubated in fresh Click-iT reaction cocktail for 30 minutes at RT in the dark. Coverslips were mounted using Prolong Gold with DAPI (Fisher Chemical).

### Antibiotic Treatment

Prior to tamoxifen treatment, 8-12 week old female and male Fmo5^F/F^ and Fmo5^IntKO^ mice underwent microbiota normalization by collecting, mixing, and redistributing the bedding of all cages involved in the experiment every other day for 2 weeks. Following microbiota normalization, mice were given IP injections of tamoxifen (100 mg/kg of body weight, MP Biochemical) for 5 days. From days 5-14 of knockout, mice were provided an antibiotic cocktail or water (control) in their water supply. The antibiotic treatment consisted of gentamycin (500 mg/L), ampicillin (1 g/L), neomycin (1 g/L), penicillin (100 U/L), and streptomycin (200 mg/L).

Mice were also given a 150 μL gavage consisting of vancomycin (1 mg/mL) and metronidazole (0.5 mg/mL) or 1x PBS (control) every other day starting on day 5 of KO. Mice were euthanized at day 14 of knockout and proximal colon samples were collected for histological analysis.

### 16s RNA sequencing

Sequencing of bacterial 16s rRNA was performed using the Microbial Systems Molecular Biology Lab which is part of the University of Michigan Host Microbiome Initiative. 8-12 week old mice were given intraperitoneal injection of 100 mg/kg tamoxifen (MP Biomedical) for 5 days.

After 14 days of KO, mice were necropsied, and fecal pellets were collected from the intestines. Using a dual index sequencing strategy, the V4 regions of the 16s rRNA was amplified. The samples were then sequenced using MiSeq Reagent kit V2 500 cycles in Illumina MiSeq.

Mothur (v1.42.3) was used to analyze bacterial communities, and quality of the samples was demonstrated via aligned reading of a DNA extraction control and genotype clustering via principal component analysis.

### RT-qPCR

RNA was extracted for gene expression analysis from colon epithelial scrapes using the chloroform method. Samples were homogenized in Trizol (Ambion), the supernatant removed, and chloroform (Sigma-Aldrich) was used to cause phase separation. Isopropanol (Acros Organics) was added to cause RNA precipitation before the pellet was washed in 70% ethanol and allowed to dry before being reconstituted with water. RNA concentrations were measured by nanodrop and synthesized into cDNA by Reverse Transcription. 1μg of RNA was loaded into each reaction including Oligo(dT) (Promega), dNTP mix (Thermo Scientific), and sterile water. Samples were heated at 65°C for 6 minutes and quick chilled on ice prior to adding First Strand Buffer (Invitrogen) and DTT (Invitrogen) and incubating at 42°C for 2 minutes. SuperScript II RT (Invitrogen) was added to each reaction and incubated at 42°C for 50 minutes prior to enzyme inactivation by heating at 70°C for 15 minutes. RT-qPCR was performed using SYBR green master mix (Applied Biosystems), sample cDNA, and forward and reverse primers (**Supplemental Table 1**). qPCR cycles were run at a standard ramp speed with each sample run in technical duplicate or triplicate. The resulting comparative ΔΔCT values were used to calculate gene expression between samples with technical replicates averaged.

### H&E staining

Samples were deparaffinized using xylene (Acros Organics) and rehydrated using washes of 100%, 95%, and 70% ethanol at room temperature. Slides were stained in Hematoxylin (Gill’s 2x) (RICCA) for 2 minutes and dipped 1-2 times in Bluing Reagent (ScyTek Laboratories). The slides were next counterstained in eosin and dehydrated using washes of 95% ethanol, 100% ethanol, and xylene. Coverslips were mounted onto slides using xylene based Permount (Fisher Chemical).

### AB/PAS staining

Samples were deparaffinized using xylene (Acros Organics) and rehydrated using washes of 100% and 95% ethanol at room temperature. Samples were incubated in 3% acetic acid for 3 minutes followed by Alcian Blue (Newcomer Supply) for 30 minutes at room temperature.

Incubation in 0.5% periodic acid (Alfa Aesar) for 10 minutes was used to oxidize samples. Slides were next incubated in Schiff’s Reagent (Electron Microscopy Sciences) at room temperature for 30 minutes. Samples were dehydrated using washes of 95% ethanol, 100% ethanol, and xylene. Coverslips were mounted onto slides using xylene based Permount (Fisher Chemical).

### Western Blot

Upon necropsy, proximal colon epithelium was harvested and flash frozen in liquid nitrogen. Protein was extracted using RIPA buffer (Thermo Fisher) with protease and phosphatase inhibitors (Thermo Fisher). Protein concentration was quantified using a Pierce BSA protein assay kit (Thermo Fisher) and normalized and samples for Western blot were prepared using RIPA buffer and loading dye to a final concentration of 4x. Samples were incubated at 95°C for 5 minutes prior to being flash frozen in liquid nitrogen and stored at -20°C until use. 6 μg of protein from each sample was loaded into a 4-20% Mini-PROTEAN® TGX gel (BioRad) and proteins were separated by size using electrophoresis. Gels were transferred onto a 0.45 µM nitrocellulose membrane by wet transfer method. Membranes were blocked in 5% milk for 1 hour at RT prior to incubation with primary antibodies (IRE1β: 1:500 and AGR2: 1:1000 from Invitrogen, and FMO5: 1:1000 from Cell Signaling) overnight at 4°C. The following day, membranes were washed and incubated in anti-rabbit secondary antibody (Cell Signaling) at a concentration of 1:2000 in 5% milk for 1 hour at RT. Proteins were visualized using an ECL kit (Thermo Fisher) and Chemidoc Imager before protein levels were analyzed using ImageJ.

### TUDCA Treatment

Mice were given IP injection of 500 mg/kg/day of Tauroursodeoxycholic Acid (TUDCA, Cayman Chemical) dissolved in 1x PBS once daily starting 7 days prior to tamoxifen treatment. TUDCA treatment was continued during the tamoxifen treatment regime, where mice received an IP injection of each chemical for 5 days. Following the conclusion of tamoxifen treatment, mice received TUDCA injections for an additional 14 days and were euthanized at 14 days of KO.

### Statistical Analyses

All data are displayed as mean +/- standard deviation. The sample size of animals in each experiment are in each figure legend, where “n” represents independent biological replicates. All plots were graphed and statistical tests were calculated using GraphPad Prism Version 10.2.2. Statistical significance for all comparisons between two experimental groups were calculated using two-tailed, unpaired t-tests. Statistical significance for all comparisons with greater than 2 experimental groups was calculated in using one-way analysis of variance (ANOVA) (or nested one-way ANOVA) with multiple comparisons and Tukey post-hoc corrections when appropriate. Statistical significance was determined for p-values less than 0.05, and are reported as * when p < 0.05, ** when p < 0.01, *** when p < 0.001, and **** when p < 0.0001. For graphs displaying multiple measurements for each independent mouse (crypt length and inner mucus barrier thickness), each plot represents the measurements from a single mouse with mean and standard deviation values for that mouse superimposed on the plot. All authors had access to study data and have reviewed, edited, and approved the final manuscript.

## ACKNOWLEDGEMENTS

We acknowledge Wanda Filipiak & Galina Gavrilina for preparation of transgenic mice in addition to the Transgenic Animal Model Core. Core support was provided by The University of Michigan Gut Peptide Research Center, NIH grant number DK34933. MLS was supported by NIH F31DK134183 and T32HD007505. SFL was supported by NIH P30AG024824 and the Glenn Foundation for Medical Research. Images were created with BioRender.com.

---

**Supplemental Figure 1.** (**A**) Relative gene expression of Fmo5 from colon epithelium of 1-, 3-, and 6-month old female and male wildtype mice (n = 4-5 mice/group). Relative gene expression of (**B**) intestinal cell type markers Tff3, Slc2a5, Alpi, Mmp7, Lys1, Pou2f3, Neurog3, ChgA, Lgr5, and Bmp4, and (**C**) absorptive and secretory progenitor cell markers Hes1 and Atoh1, respectively, from colon epithelium of ∼10 week old female (F) and male (M) Fmo5^F/F^ and Fmo5^IntKO^ mice (n = 3-6 mice/group). Plots display mean +/- SD for each group. Statistical significance was calculated by 1-way ANOVA with Tukey correction for multiple comparisons. * p < 0.05.

**Supplemental Figure 2.** (**A**) Relative gene expression of canonical ER stress markers BIP, CHOP, IRE1α, PERK, ATF6, and ATF4 from colonic epithelium of female Fmo5^F/F^ and Fmo5^IntKO^ mice 14 days after tamoxifen treatment (n = 4-5 mice/group). (**B**) Representative Western blots of BIP and PERK proteins with β-actin controls from female Fmo5^F/F^ and Fmo5^IntKO^ mice at 14 days of KO. (**C**) Quantification of BIP and PERK protein levels described in **B**, normalized to their respective β-actin levels (n = 3-6 mice/group). (**D**) Relative gene expression of Spliced- and Total- XBP1 in colon scrapes from female Fmo5^F/F^ and Fmo5^IntKO^ mice 14 days post-tamoxifen treatment (n = 4-5 mice/group). The ratio of Spliced XBP1 to Total XBP1 was calculated for each mouse. Plots display mean +/- SD for each group. Statistical significance was calculated by unpaired t-test.

