## Supplemental Figures for "Fmo5 plays a sex-specific role in goblet cell maturation and mucus barrier formation"

### Supplemental Figure 1.

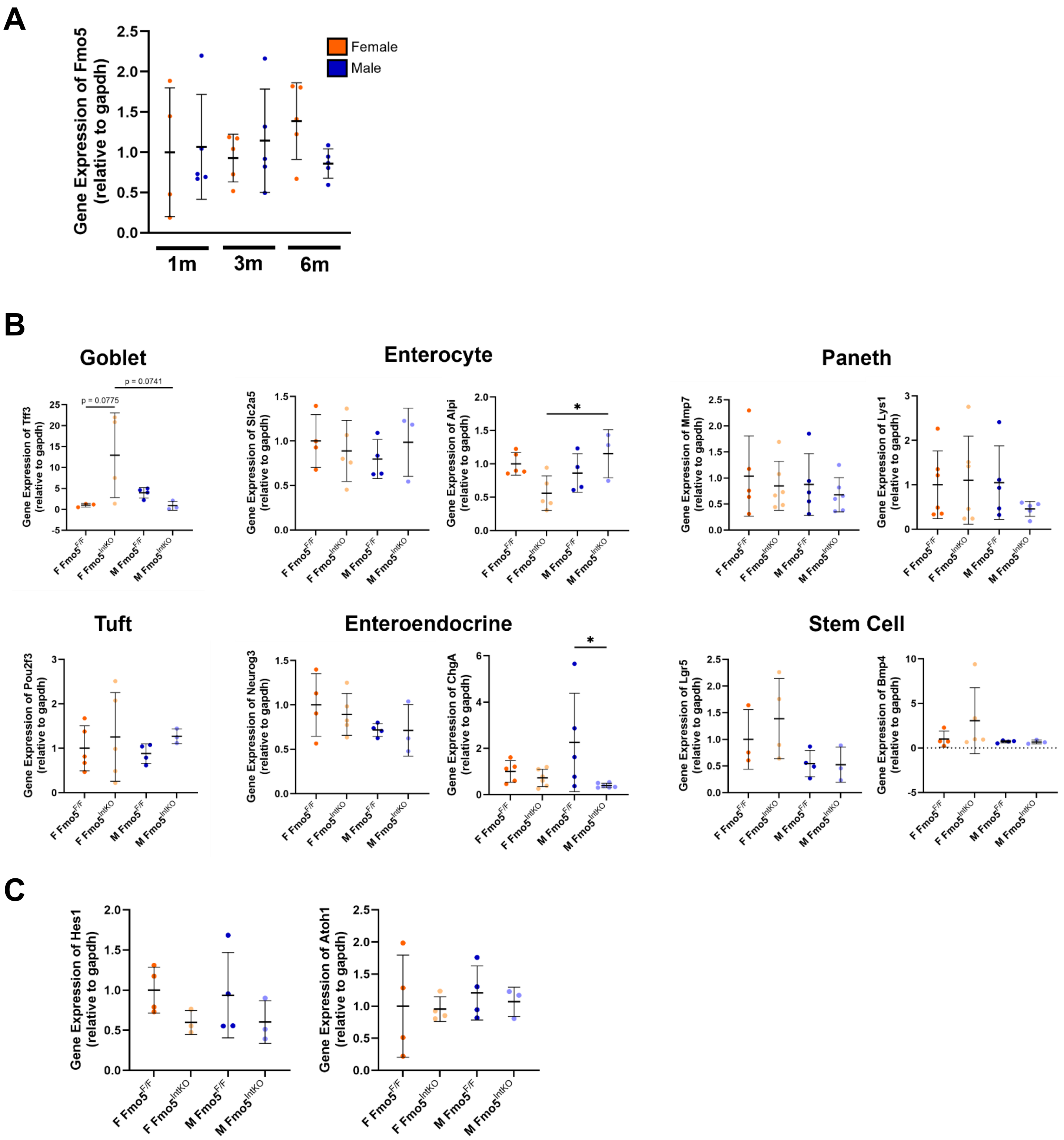


**Supplemental Figure 1.** (**A**) Relative gene expression of Fmo5 from colon epithelium of 1-, 3-, and 6-month old female and male wildtype mice (n = 4-5 mice/group). Relative gene expression of (**B**) intestinal cell type markers Tff3, Slc2a5, Alpi, Mmp7, Lys1, Pou2f3, Neurog3, ChgA, Lgr5, and Bmp4, and (**C**) absorptive and secretory progenitor cell markers Hes1 and Atoh1, respectively, from colon epithelium of ~10 week old female (F) and male (M) Fmo5$^{F/F}$ and Fmo5$^{IntKO}$ mice (n = 3-6 mice/group). Plots display mean +/- SD for each group. Statistical significance was calculated by 1-way ANOVA with Tukey correction for multiple comparisons. * $p < 0.05$.

### Supplemental Figure 2.

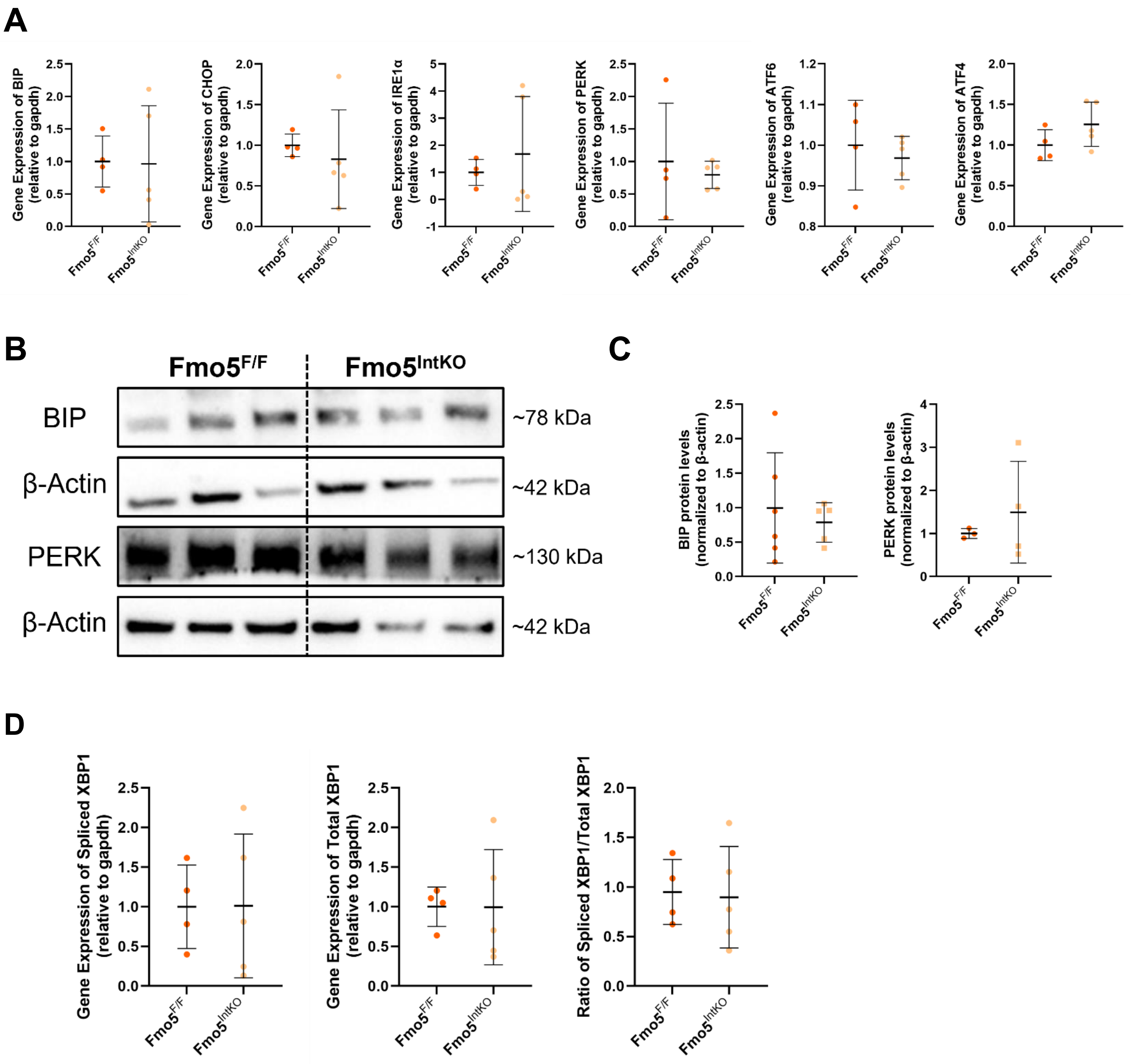


**Supplemental Figure 2.** (**A**) Relative gene expression of canonical ER stress markers BIP, CHOP, IRE1α, PERK, ATF6, and ATF4 from colonic epithelium of female Fmo5$^{F/F}$ and Fmo5$^{IntKO}$ mice 14 days after tamoxifen treatment (n = 4-5 mice/group). (**B**) Representative Western blots of BIP and PERK proteins with β-actin controls from female Fmo5$^{F/F}$ and Fmo5$^{IntKO}$ mice at 14 days of KO. (**C**) Quantification of BIP and PERK protein levels described in **B**, normalized to their respective β-actin levels (n = 3-6 mice/group). (**D**) Relative gene expression of Spliced- and Total- XBP1 in colon scrapes from female Fmo5$^{F/F}$ and Fmo5$^{IntKO}$ mice 14 days post-tamoxifen treatment (n = 4-5 mice/group). The ratio of Spliced XBP1 to Total XBP1 was calculated for each mouse. Plots display mean +/- SD for each group. Statistical significance was calculated by unpaired t-test.

### Supplemental Table 1.

| qPCR Primer Sequences | | |
|---|---|---|
| **Gene** | **Forward Primer** | **Reverse Primer** |
| **Fmo5** | 5'-TCCGAACTAGTGTAGCAGCG- 3' | 5'-CAGGCCACAGAAAAGATAACTTCG- 3' |
| **Muc2** | 5'-GCTGACGAGTGGTTGGTGAATG- 3' | 5'-GATGAGGTGGCAGACAGGAGAC- 3' |
| **Tff3** | 5'-GCACCATACATTGGCTTGG- 3' | 5'-AGAGCCCTCTGGCTAATGCT- 3' |
| **Slc2a5** | 5'-GCGATTCCTACTCCTCGTCG- 3' | 5'-CGTCAGCACTAAGCAGGCTA- 3' |
| **Alpi** | 5'-CAGAACCTGGTGCAAACGTG- 3' | 5'-TTCCAAACATACCGGGCTCC- 3' |
| **Mmp7** | 5'-CAGACTTACCTCGGATCGTAGTGG- 3' | 5'-GTTCACTCCTGCGTCCTCACC- 3' |
| **Lys1** | 5'-GAATGCCTTGGGGATCTCTC- 3' | 5'-CTGTGGGATCAATTGCAGTG- 3' |
| **Pou2f3** | 5'-GGTGAATCTGGAGCCCATGC- 3' | 5'-GGTGTCCGTGGAATCAGCGA- 3' |
| **Neurog3** | 5'-TACCTCACTGGCTCCTCCAT- 3' | 5'-TGTAGCGGGCAGTAAAGACG- 3' |
| **ChgA** | 5'-GTCTCCAGACACTCAGGGCT- 3' | 5'-ATGACAAAAGGGGACACCAA- 3' |
| **Lgr5** | 5'- CGGAGGAAGCGCTACAGA - 3' | 5'- CTGGGTGGCACGTAGCTG - 3' |
| **Bmp4** | 5'-TGAGTACCCGGAGCGTCC- 3' | 5'-CTCCAGATGTTCTTCGTGATGG- 3' |
| **Hes1** | 5'-AGTGTCACCTTCCAGTGGCT- 3' | 5'-TGGGCTAGGGACTTTACGGG- 3' |
| **Atoh1** | 5'-GCCTTGCCGGACTCGCTTCTC- 3' | 5'-TCTGTGCCATCATCGCTGTTAGGG- 3' |
| **BIP** | 5'-GGCCGAGACAACACTGACCT- 3' | 5'-CGACGGTTCTGGTCTCACACA- 3' |
| **CHOP** | 5'-TATCTTGAGCGTAACACGTCGAT- 3' | 5'-TGGAACACTCTGCTCAGGT- 3' |
| **IRE1α** | 5'-TGCTGCAGCTTGTGACGTTG- 3' | 5'-AGTGCATGGAGACTTCCGTCC- 3' |
| **PERK** | 5'-TTTTCCATCCTGAGCCCCACA- 3' | 5'-TCGGCACTCAGGGAGTCGTA- 3' |
| **ATF6** | 5'-ACCTGTGGCTTGTGGGTGTTT- 3' | 5'-TTGGTCCATCGTGGGAGGACA- 3' |
| **ATF4** | 5'-GAGCACAGCGGAGAAGGGTT- 3' | 5'-GGTCACGTGATCCTACCGCA- 3' |
| **Spliced XBP1** | 5'-CTGAGTCCGAATCAGGTGCAG- 3' | 5'-GTCCATGGGAAGATGTTCTGG- 3' |
| **Total XBP1** | 5'-TGGCCGGGTCTGCTGAGTCCG- 3' | 5'-GTCCATGGGAAGATGTTCTGG- 3' |
| **Gapdh** | 5'-ACTGAGCAAGAGAGGCCCTA- 3' | 5'-TATGGGGGTCTGGGATGGAA- 3' |
